# A highly-selective chloride microelectrode based on a mercuracarborand anion carrier

**DOI:** 10.1101/629691

**Authors:** Marino DiFranco, Marbella Quinonez, Rafal M. Dziedzic, Alexander M. Spokoyny, Stephen C. Cannon

## Abstract

The chloride gradient plays an important role in regulating cell volume, membrane potential, pH, secretion, and the reversal potential of inhibitory GABA_A_ receptors. Measurement of intracellular chloride activity, 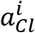, using liquid membrane ion-selective microelectrodes (ISM), however, has been limited by the physiochemical properties of Cl ionophores which have caused poor stability, drift, sluggish response times, and interference from other biologically relevant anions. Most importantly, intracellular HCO_3_^−^ may be up to 4 times more abundant than Cl^−^ (e.g. skeletal muscle) which places severe constraints on the required selectivity of a Cl – sensing ISM.

Previously, a sensitive and highly-selective Cl sensor was developed in a polymeric membrane electrode (Badr et al. 1999) using a trinuclear Hg(II) complex containing carborane-based ligands, [9]-mercuracarborand-3, or MC3 for short. Here, we have adapted the use of the MC3 anion carrier in a liquid membrane ion-selective microelectrode and show the MC3-ISM has a linear Nernstian response over a wide range of *a_Cl_* (0.1 mM to 100 mM), is highly selective for Cl over other biological anions or inhibitors of Cl transport, and has a response time of less than 5 sec. Importantly, over the physiological range of *a_Cl_* (1 mM to 100 mM) the potentiometric response of the MC3-ISM is insensitive to HCO_3_^−^ or changes in pH. Finally, we demonstrate the biological application of an MC3-ISM by measuring intracellular *a_Cl_*, and the response to an external Cl-free challenge, for an isolated skeletal muscle fiber.

## INTRODUCTION

Chloride ions (Cl^−^) are the major inorganic free anions of the extracellular fluid, and together with organic bicarbonate ions (HCO_3_^−^) are the major intracellular anions [29,32]. The transmembrane Cl^−^ gradient has a critical role in multiple cellular processes including cell volume regulation, acid-base homeostasis, fluid and electrolyte secretion, membrane excitability and the reversal potential of GABA_A_-receptors at inhibitory synapses (see [29] and references therein). Moreover, the Cl^−^ gradient is dynamic in many cells. For example, a developmental change in the expression of cation / chloride co-transporters reduces [Cl^−^]_i_ in mature neurons, and thereby converts GABA_A_-receptor mediated responses from excitatory to inhibitory by shifting *E_Cl_* to more negative potentials [27]. The steady-state Cl^−^ gradient is determined by the balance of both passive transport, through ion channels as determined by the Cl^−^ electrochemical potential, and by secondary active mechanisms via coupling to exchangers or cotransporters.

We sought a method to measure the Cl^−^ gradient across the sarcolemma of skeletal muscle, where the resting potential (*V_rest_*) is strongly dependent on the Cl^−^ driving force, *V_m_* – *E_Cl_*, because these cells have a remarkably high resting chloride conductance (*g_Cl_* ~1 mS/cm^2^) [14,26]. Moreover, shifts of the sarcolemmal Cl^−^ gradient have been causally implicated in attacks of periodic paralysis [23,37]. Understanding the roles of Cl^−^ in muscle physiology, and all cells in general, demands knowing its concentration in the intra- and extracellular milieus, or more properly its activity (*a_Cl_*) or “effective concentration”, that ultimately determines the physiochemical reactivity of Cl^−^ [33].

While measuring the extracellular 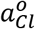 is relatively easy, non-destructive measurements of intracellular 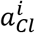 poses major challenges. Basically, two methods are available to assess the intracellular chloride activity: (1) micro-photometric techniques, using either chemical or optogenetic Cl-sensors [13,3] and (2) potentiometric techniques with liquid membrane ion-selective micro-electrodes (ISM for short) [31,18]. Photometric techniques allow for specific, high time resolution and space-resolved determination of changes in the activity of physiologically relevant ions, e.g. Ca^2+^, Na^+^, K^+^, H^+^, Cl^−^. However, activity-dependent changes in optical properties of sensors are usually difficult to de-convolve into accurate molar units, and in some conditions, sensors can alter the extent and kinetics of the ion concentration changes aimed to be measured. Fluorescence life-time imaging circumvents some of these challenges [13], but requires specialized and expensive detection systems. In principle, an ISM is devoid of these limitations, and is expected to provide a direct determination of intracellular activity. In fact, ISMs are often used to measure the activities of ions in solutions that are used to calibrate photometric sensors. On the other hand, ISMs cannot track fast changes in ion activity (i.e. in the ms range, but see [12,21]) and in some cases selectivity is a limiting factor. The key component of an ISM is the ion-selective carrier or ionophore, which endows the electrode with the selectivity required to discriminate between ions of similar nature and the sensitivity (usually down to the sub-micromolar range). In contrast to the case for cations, there is a general lack of highly-selective naturally occurring or synthetic anion ionophores, and in particular for Cl^−^ [2,10].

We tested ISMs made with several commercially available Cl-ionophores, and unlike our experience with cation ionophores (H^+^, Na^+^) we found their responses far from ideal. These Cl-ionophores included: tributyl tin (TBT), chloride ionophores I-IV (Selectophore™, Millipore-Sigma), and the antibiotic 3,4,4’-trichlorocarbanilide [28]. We found ISMs fabricated with these compounds had one or more of the following problems: lack of linearity, sub- or supra-Nernstian responses, drift, hysteresis, poor solubility in liquid membranes, sensitivity to blockers of chloride channels or transporters, and poor selectivity for Cl^−^ over HCO_3_^−^. We also tested a chemically diverse group of recently published compounds described as Cl^−^ carriers for biological membranes: compound-1 [30], compound 4H [9], cholapod-3, and decalin-13 [10,19,20], and found them not appropriate to build ISMs, mostly demonstrating solubility problems.

Here we report that a Cl-selective anion carrier based on a trinuclear Hg(II) complex containing carborane-based ligands, [9]mercuracarborand-3 (MC3), that was previously used in polymer membrane Cl-sensors [5] is also ideal for liquid membrane Cl-selective ISMs. MC3 is a macrocyclic structure bearing a pre-organized Lewis acid cavity that selectively complexes with Cl^−^ and functions as an anion carrier in the liquid membrane ISM. We show the MC3-ISM is highly selective for Cl^−^ over other biologically relevant anions (HCO_3_^−^, lactate, PO_4_^−^) and is insensitive to Cl-channel blockers (9-anthracene carboxylic acid), inhibitors of cation-chloride co-transporters (bumetanide), or changes in pH and is therefore well-suited for biophysical applications to quantitatively and accurately measure 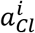.

## METHODS

### MC3 synthesis

[9]mercuracarborand-3 (MC3) was synthesized using air-free Schlenk techniques as described by Hawthorne and co-workers [38]. An oven dried Schlenk flask was charged with 144 mg (0.5 mmol) of *ortho*-carborane (Boron Specialties), filled with an N_2_ atmosphere, and 5 mL of anhydrous diethyl ether (Et_2_O) (Fisher, inhibitor free) were added via syringe. The reaction vessel was cooled to 0 °C using an ice bath, 0.85 mL of 2.5 M n-BuLi in hexane (Sigma-Aldrich) was added dropwise to the rapidly stirring reaction mixture. After addition of the n-BuLi solution, the ice bath was removed and the reaction was stirred at room temperature for 2 hours. Next, 315 mg of Hg(OAc)_2_ (Mallinckrodt) was added to the reaction mixture under a flow of N_2_ and the reaction mixture was stirred at room temperature for 16 hours. Upon completion of the reaction, the reaction was quenched with 5 mL of H_2_O, the organic layer was collected, the aqueous portion was extracted with 5 mL Et_2_O. The organic portions were combined and washed with 5 mL H_2_O, then the organic layer was dried with MgSO_4_ and filtered through 2 cm of silica gel using Et_2_O as the eluent. The filtrate was evaporated to dryness using a rotary evaporator to yield a white solid. Due to difficulty with purifying the product by recrystallization from Et_2_O, an alternative purification method was developed as follows. The white solid was washed with hexanes, dissolved in 10 mL of 20% (v/v) hexanes in Et_2_O and left in a fume hood to evaporate the Et_2_O portion which resulted in precipitation of an off-white powder. The supernatant was decanted and the solids were loaded onto a 3 cm plug of silica gel wetted with 40% acetone (v/v) in hexanes and eluted with 40% acetone (v/v) in hexanes. The resulting filtrate was evaporated to dryness to yield 50 mg (14% yield) of an off-white solid which contains >95% MC3 by ^11^B NMR. ^1^H NMR (400 MHz, (CD_3_)_2_CO) δ 3.0-1.2 (m). ^11^B NMR (160 MHz, (CD_3_)_2_CO) δ 1.6 (d, 2B), −5.4 (d, 2B), −8.0 (d, 6B).

### Micro-pipette silanization

Micro-pipettes were pulled from 1.5 mm borosilicate capillaries with micro-filaments (BF150-86-10, Sutter Instruments). A 4-step pulling protocol was optimized to obtain a short taper (~8 mm) leading to a sub-micron tip opening, using a horizontal puller (P97, Sutter Instruments). When filled with a saline mimicking the ion composition of myoplasm (see solutions), the microelectrodes had tip resistances of 10-12 MΩ. The use of capillaries with micro-filaments greatly reduced the filling-time in the manufacture of ISMs, without compromising the sealing of the organic phase to the inner surface of the capillary wall. Capillaries were silanized as supplied from the vendor, with no pre-treatment.

For silanization, micro-pipettes were laid horizontally on a support in a Pyrex glass jar (with lid, ~500 ml). One hundred μl of chlorotrimethylsilane (Sigma) was placed in the jar and allowed to evaporate at RT for ~5 min. The silane vapor had free access to a micro-pipette’s interior and exterior surfaces. Then, the micro-pipettes were baked at 250°C for at least 4h (usually overnight). As occurs with all liquid-junction microelectrodes, silanization was essential to insure the mechanical stability and high electrical resistance between the hydrophobic liquid membrane and the inner wall of an ISM. Silanized micro-pipettes were stored dry in a closed jar (WPI Jar-E215) to avoid dust adhesion, and were used for up 2 weeks with no noticeable deterioration.

### Cl sensing liquid membrane and ISM half-cell saline

The ISM liquid membrane consisted of a short column (200-300 μm) of a hydrophobic “chloride cocktail” lodged in the tip of the micropipette. The chloride cocktail was made by dissolving the MC3 and a lipophilic cationic additive (tridodecylmethyl ammonium chloride, TDMAC, Sigma) in the water-immiscible organic solvent 2-nitrophenyl octyl ether (NPOE, Sigma). We used 10% MC3 and 2.5% TDMAC in NPOE (w/w). No other mixing ratios or solvents were studied. The ISM half-cell consisted of a Cl-based saline, backfilled into the microelectrode and in contact with an Ag/AgCl reversible electrode.

### ISM fabrication

We devised a simple and efficient method to fabricate ISMs. Starting with previously silanized micro-pipettes, we routinely made six ISMs in less than 1 h. To fabricate an MC3-ISM both the chloride cocktail and the reference saline were sequentially backfilled into the micro-pipette, while viewing under a stereomicroscope. First, an excess amount of chloride cocktail (2-3 times the final volume of the liquid membrane) was delivered as close as possible to the micro-pipette tip using a 34G microneedle (Quickfil 34G-5, WPI, for which the polyimide coating was stripped from the end). The organic phase spontaneously moves towards the tip of the micro-pipette by capillary action and fills the shaft in 3-5 min. Initially using excess chloride cocktail speeds up this filling step. Once the tip is filled, the chloride cocktail volume is reduced as much as possible by suction with a new 34G needle. Finally, the saline solution is delivered (using another 34G needle), assuring that a clean interface is formed between the two liquids. Care must be taken to avoid air bubbles in the organic phase or interface, otherwise the ISM should be discarded.

Prior to use, each MC3-ISM was individually tested using a 3 point calibration including solutions of 1, 10 and 100 mM Cl (see solutions), with *a_Cl_* of 0.64, 6.46, and 68.5 mM respectively. Tested MC3-ISMs were stored in closed jars (to avoid evaporation and dust) with tips submersed in a filtered (0.2 μm) solution containing 10 mM NaCl; and could be used after 1-2 weeks from fabrication with no detectable deterioration.

### Measurement of ISM potential

A two-channel high input impedance (>10^15^ Ω) amplifier (FD223a, WPI) was used to measure the electrical potential of MC3-ISMs. One channel was connected to the Ag/AgCl wire of an MC3-ISM. The second channel was connected to a standard (“sharp”) microelectrode having 10-15 MΩ tip resistance when filled with electrode solution (see solutions). This microelectrode served as a reference electrode, whose value (V_ref_) reflects changes in junction potential that may occur upon solution exchanges. Three amplifier outputs, V_ref_, V_ISM_, and the analog difference V_ISM_ – V_ref_ were digitized with a 16-bit A/D converter and stored on a computer (LabVIEW, National Instruments). The figures show V_Cl_, defined as – (V_ISM_ – V_ref_), because with this sign convention the potentiometric signal increases for an increase in *a_Cl_*.

### Solutions

Solutions containing different [Cl^−^] were prepared in two ways: (1) Calibration solutions devoid of any possible interfering anion were prepared by serially diluting a pure NaCl solution with distilled water (80 MΩ, Millipore). (2) Calibration solutions suitable for use with mammalian cells were prepared by mixing two solutions having a composition similar to that of Tyrode’s buffer but differing in the main anion content, and having the same pH and divalent cation content. These solutions differ in that one (150-Cl solution) contained Cl^−^ as the main anion, and the other (0-Cl solution) was devoid of Cl^−^, and contained SO_4_^-2^ as the main anion. Both solutions have HEPES (a possible interfering anion) as the pH buffer.

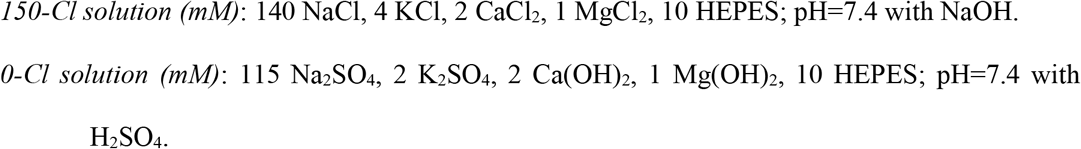

Calibration solutions containing from 100 mM to 10 nM nominal Cl concentrations were made by serial dilutions of the 150-Cl solution with the 0-Cl solution.

Solutions containing potential interfering anions were prepared by adding 10 mM of Na-lactate, NaHCO_3_, Na_2_HPO_4_ or NaSCN to a mixture of the 150-Cl and 0-Cl solutions, and then performing a similar serial dilution as described above to obtain the desired [Cl^−^]. Drug-containing solutions were made from highly concentrated stock solutions of 9-anthracenecarboxylic acid (9-ACA, 500 mM in DMSO) or bumetanide (BMT, 50 mM in DMSO). DMSO final concentration was kept below 1:1000, and controls were run to exclude direct effects of solvent.

Solutions used for recording from isolated mouse muscle fibers were either standard (i.e. Cl-containing) Tyrode’s solution or Cl-free Tyrode’s in which the Cl^−^ was replaced with SO_4_^−^. Standard Tyrode’s (mM): 150 NaCl, 4 KCl, 2 CaCl_2_, 1 MgCl_2_, 10 glucose, 10 HEPES; pH=7.4 with NaOH. Cl-free Tyrode’s (mM): 115 Na_2_SO_4_, 2 K_2_SO_4_, 2 Ca(OH)_2_, 1 Mg(OH)_2_, 10 HEPES; pH=7.4 with H_2_SO_4_. An agar bridge was used to connect the bath of the recording chamber to the reference ground, so that shifts in the junction potential were minimized when bath exchanges were used.

The MC3-ISM was backfilled with a solution containing (in mM): 100 NaCl, 10 HEPES, pH=7.4 with H_2_SO_4_. The reference (bath) and intracellular microelectrodes were filled with a solution (in mM): 70 K-aspartate, 5 di-Na ATP, 5 di-tris creatine phosphate, 40 EGTA, 20 MOPS, 10 MgCl_2_, pH=7.4 with KOH.

All chemicals were purchased and used as received from Millipore-Sigma, unless otherwise noted (*vida supra*).

### Cl activity calculation (*a_Cl_*)

The value for the activity of an ion in solution, *a_ion_* is often obtained from tables [25], but these values were calculated from solutions of single salts (e.g. NaCl, KCl). Instead, we calculated *a_Cl_* (in mM) for all simple and mixed solutions as: *a_Cl_* = *γ_Cl_* * [*Cl*], where *γ_Cl_* is the chloride activity coefficient (dimensionless). The activity coefficient was calculated from the Debey-Hückel [11] formalism as:

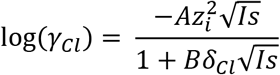

where *δ_Cl_* is the effective Cl^−^ ion diameter (0.3 nm), *A* = −0.510, and *B* = 3.29. The ionic strength, *Is*, of a solution containing many ions, *ion_i_*, with valences *Z_i_* was calculated as:

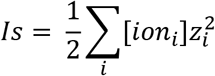

### “In cuvette” MC3-ISM characterization

A small circular plastic experimental chamber (8 mm in diameter, 1 mm depth, ~0.5 ml in volume) was used to study the response of MC3-ISMs to [Cl^−^] changes and to determine selectivity. The chamber was continuously perfused with the test solution of choice and the entire chamber could be completely exchanged in ~1 sec. Perfusion flux (up to 1-2 ml/sec) was driven by gravity and the efflux was removed by aspiration, keeping the level of solution constant to minimize drifts in junction potentials. The chamber solution was connected to ground via a salt bridge (3% agar in 125 mM KCl, w/v containing a floating platinum wire) in contact with a 125 mM KCl solution and an Ag/AgCl electrode. This arrangement represented the second (electrical) half-cell across the MC3-ISM’s liquid membrane.

### Muscle fiber ISM measurements

To test the MC3-ISM in a cellular context, we used enzymatically dissociated skeletal muscle fibers from the flexor digitorum brevis (FDB) of the mouse, as previously described [36]. All mouse procedures were approved by the University of California Los Angles Institutional Animal Care and Use Committee. Fibers were plated in the same chamber as above, and impaled with a MC3-ISM and a reference microelectrode. Before impalement, both MC3-ISM and reference micro-electrode were zeroed in the presence of 100-Cl Tyrode.

### Selectivity coefficient, 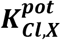

The relative selectivity of an MC3-ISM for Cl^−^ over a test interfering anion was determined by the fixed interference method [34]. The concentration of the test interfering anion, [X^−^], was maintained at a fixed value of 10 mM, and the MC3-ISM potential was measured as the [Cl^−^] was varied from 10 nM to 100 mM. The MC3-ISM potential was plotted against log(*a_Cl_*) and the intersection of the two linear portions (Nernstian for high [Cl^−^] and saturating for low [Cl^−^]) determined the value of *a_Cl_* that was used to calculate the selectivity coefficient for Cl^−^ over X^−^ as:

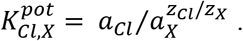

Values are reported as the mean ± standard error of the mean.

## RESULTS

### A cationic additive increases the Cl-responsiveness an MC3-based ISM

We first tested the response of liquid membrane ISMs constructed using a cocktail containing only MC3 (10%) and NPOE. These electrodes were only weakly sensitive to [Cl^−^] changes, as shown for an exemplary response in Figure 1A. This response is sub-Nernstian, with a slope of 9.2 ± 0.67 mV/decade (n = 3) on a plot of V_Cl_ as a function of log_10_(*a_Cl_*) (Figure 1C, blue symbols). This result is in agreement with previous data [5], demonstrating that polymeric membranes containing MC3 are insensitive to Cl unless the membranes are doped with cationic compounds.

**Fig. 1.**
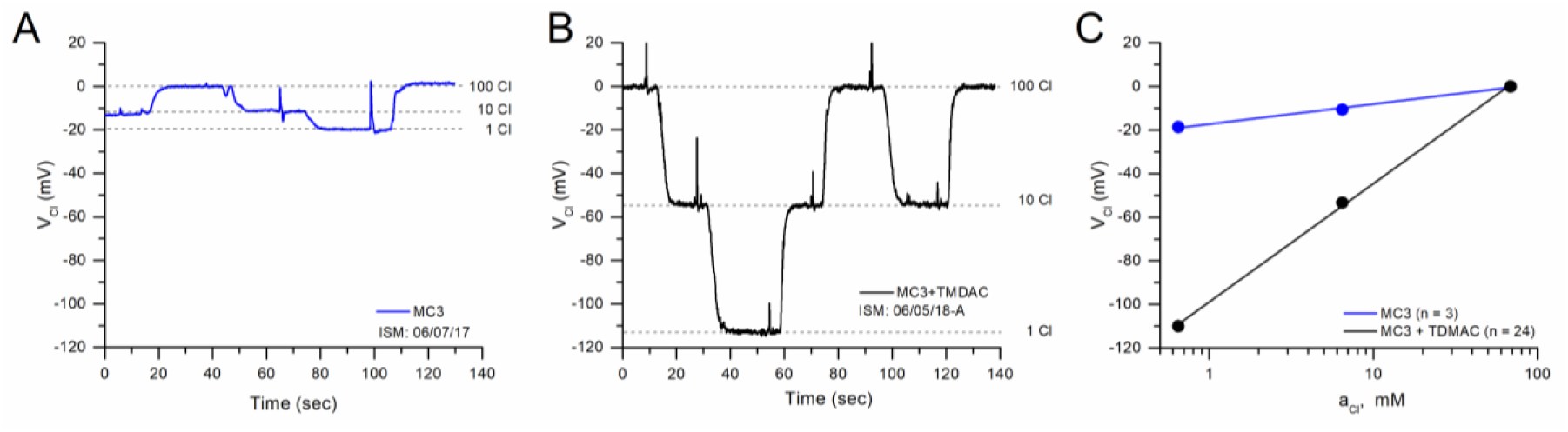
TDMAC is required for Nernstian behavior of the MC3-ISM. (A). Response of an exemplary ISM for which the liquid membrane consisted only of 10% MC3 dissolved in NPOE, as the [Cl^−^] was varied from 1 to 100 mM. (B) Response of another ISM containing the same liquid membrane as in A, but supplemented with 2.5% TDMAC. The rapid transients for the tracings in (A) and (B) were caused by solution switching and suction-dependent fluctuations. (C) Plot of V_Cl_ as a function of log_10_(*a_Cl_*) shows a linear relation with a slope of 54.6 ± 0.18 mV (n = 24) for the ISMs containing TDMAC (black circles), as expected for a Nernstian response. For ISMs lacking TDMAC (blue circles), a sub-Nernstian response was observed with a slope of 9.2 ± 0.67 mV (n = 3). The error bars for the SEM are very small, and obscured by the symbols.

When the liquid membrane cocktail was supplemented with a lipophilic cationic additive, TDMAC (2.5%, w/w), the potentiometric response became highly sensitive to [Cl^−^] changes between 1 and 100 mM (Figure 1B). For brevity, we refer to microelectrodes with this complete liquid membrane cocktail as MC3-ISMs, but it should be recognized they are doped with TDMAC. The voltage response of MC3-ISMs was stable and showed no signs of hysteresis. Over this range of [Cl^−^], the MC3-ISM response was Nernstian, with a slope of 54.6 ± 0.18 mV/decade (n = 24), as shown in Figure 1C (black symbols). The reference electrode had very small voltage changes in response to these solution exchanges (2-3 mV, not shown), and these small deviations were subtracted from V_ISM_ to obtain V_Cl_ values shown in Figure 1.

We did not explore other combinations of MC3 and TDMAC since the 10% and 2.5% (w/w), respectively, mixture tested gave optimal results. The need for a lipophilic cation stems from the fact that MC3 is a neutral carrier, and as such it is expected to produce a limited anion response unless a counterion for the mercuracarborand complex is added [5].

### Linearity and Cl^−^ detection limit of the MC3-ISM

The linear range for Cl^−^ responsiveness of the MC3-ISM was determined by changing the [Cl^−^] over 7 orders of magnitude, in ten-fold steps from 100 mM to 10 nM. This calibration was performed with solutions prepared by serial dilution of 100 mM NaCl with water (Methods) to minimize the possibility of interfering anions at low concentrations of Cl^−^. A typical continuous recording of V_Cl_ during the decrease and then increase of [Cl^−^] is shown in Figure 2A. Jumps of V_Cl_ to new stable values are clearly evident with each solution exchange, and the dotted lines emphasize that no hysteresis was detected. The change in V_Cl_ became compressed (i.e. smaller jumps) for test solutions of less than 0.1 mM [Cl^−^], as the linear range was exceeded.

**Fig. 2.**
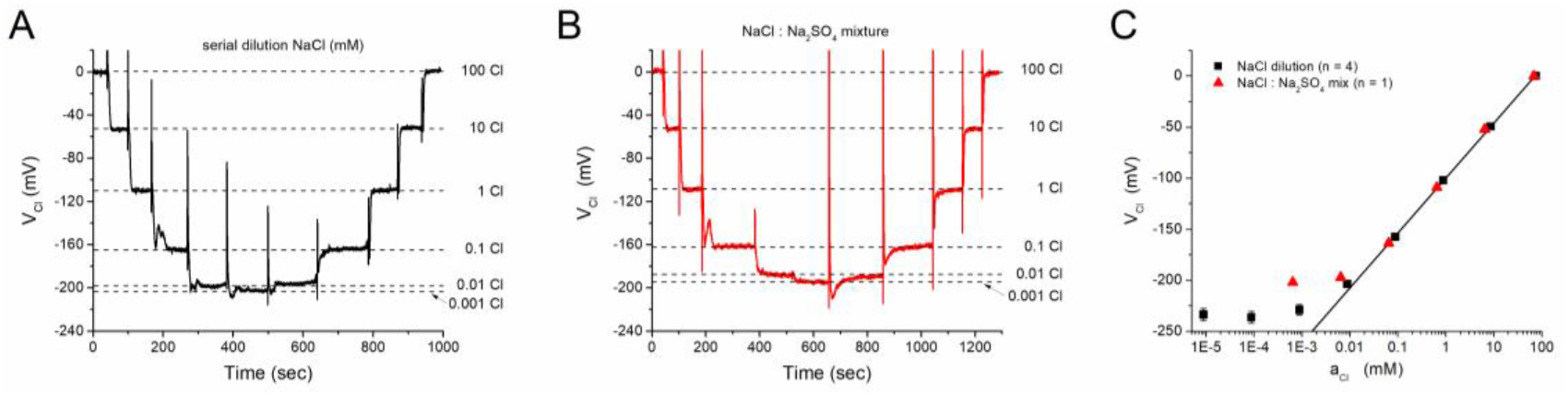
MC3-ISMs have a linear response to [Cl^−^] changes from 0.1 to 100 mM. (A) Continuous record of the response (V_Cl_) for an MC3-ISM exposed to solutions of various [Cl^−^] from 0.001 mM to 100 mM, obtained by serial dilutions of a 100 mM NaCl solution with water. (B) Continuous record of another MC3-ISM exposed to the same nominal [Cl^−^], but obtained by mixing Cl-based and SO_4_-based solutions (see Methods). The dashed lines in A and B represent the steady state values of the ISMs at the indicated [Cl^−^]. (C) The steady-state V_Cl_ is plotted as a function of *a_Cl_*, which was calculated on the basis of the ionic strength of the test solutions (See Methods). A linear fit of V_Cl_ as a function of *a_Cl_* over a range from 0.088 to 75 mM, for the dilution method, had a slope of 52.8 ± 0.63 mV/decade (n = 4, straight line)

A limited number of trials were also performed with solutions prepared by serially diluting 100 mM NaCl with a Na_2_SO_4_-based solution (Figure 2B) that is commonly used for “chloride-free” conditions with mammalian cells (Methods, 0 – Cl solution). This alternative method maintains a high ionic strength for all test values of [Cl^−^] and provides a check for evidence of interfering anions in a mammalian physiologic buffer. A plot of steady-state V_Cl_ as a function of log_10_(*a_Cl_*) shows that between 0.1 mM and 100 mM the response was linear (Figure 2C). This range exceeds the expected values of intracellular *a_Cl_* to be encountered in mammalian cells (3 to 20 mM). Moreover, the response for dilutions with pure water was near-linear down to an activity of 0.01 mM. The saturation of V_Cl_ at about −200 mV for the NaCl:Na_2_SO_4_ mixture reveals an interfering anion effect of SO_4_^-2^, although this was detectable only at low *a_Cl_* levels (< 0.1 mM) that are not encountered in cells.

The average slope of the linear response range was 52.8 ± 0.63 mV/decade (n = 14). While this slope is slightly smaller than the theoretical value of 58 mV, it is comparable to values reported for MC3 Cl-electrodes fabricated in solid membranes [5]. Those electrodes also showed a sub-Nernstian response, with the slope being smaller for lower molar ratios of TDMAC to MC3.

### The settling time for an MC3-ISM is a few seconds

Another desirable feature of ISMs for biological applications is a small time constant. To assess the time response of the MC3-ISM, we increased the flow rate of the solutions perfusing the experimental chamber, as compared with those used in experiments in Figures 1 and 2. Photometric (fluorescence) measurements (not shown) demonstrated that the contents of the experimental chamber (~0.5 ml) can be exchanged in ~1 sec. The MC3-ISM was exposed to short pulses (~ 50 sec) of 100, 10 and 1 mM NaCl. The potentiometric response, V_Cl_, reached steady state in less than 5 sec and was approximated well by a single exponential (Figure 3). The time constant for high to low [Cl^−^] transitions were about 2.5 s, whereas for transitions from low to high [Cl^−^] it was about 1.5 s. This difference was consistent amongst several different electrodes and likely reflects flow-dependent properties of the chamber, wherein an increase of [Cl^−^] occurs more abruptly than with washing out Cl^−^ from a disk-shaped chamber.

**Fig. 3.**
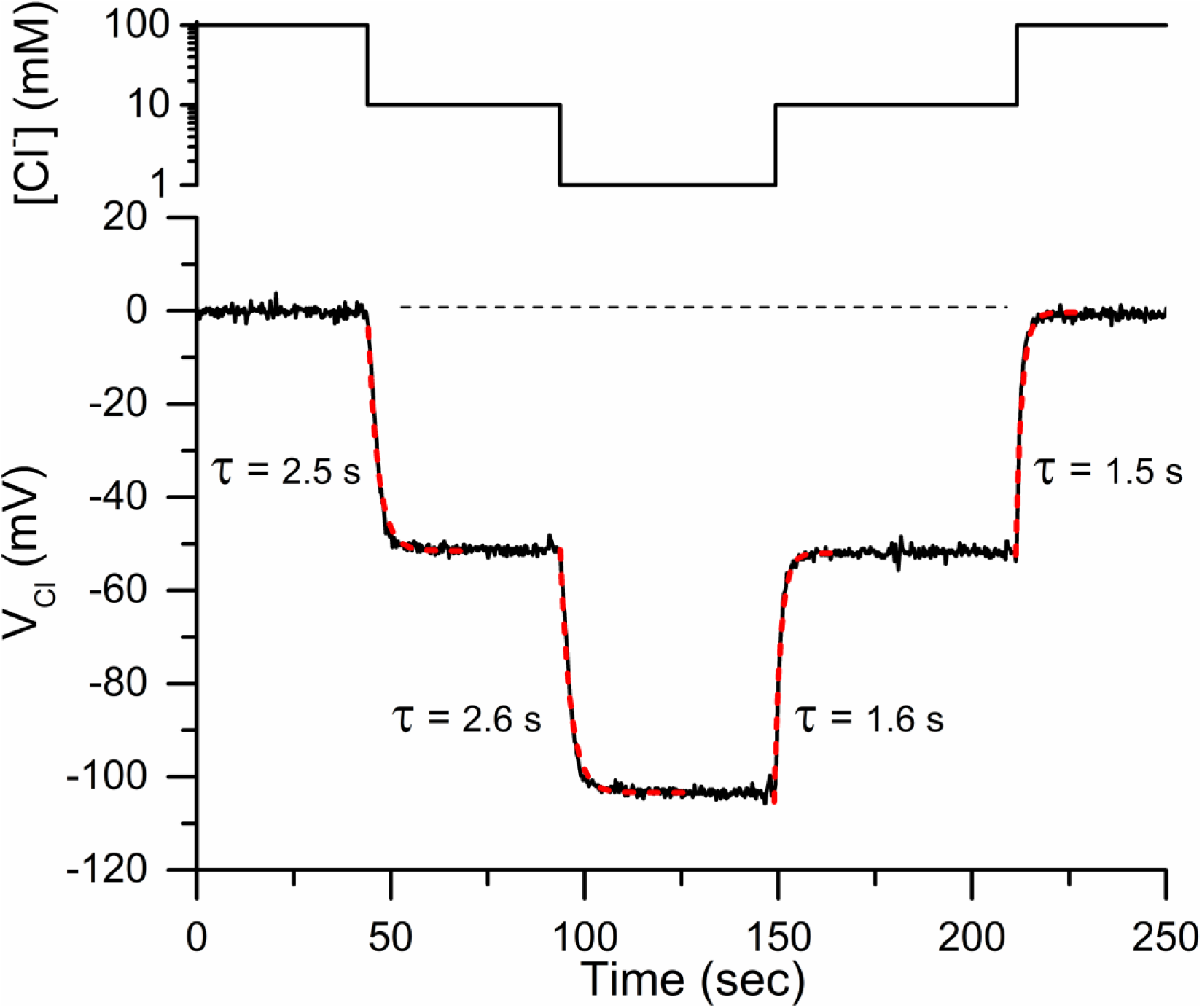
Liquid-membrane MC3-ISMs respond rapidly to [Cl^−^] changes. A continuous record of V_Cl_ (bottom) is shown in response to fast changes of [Cl^−^] between 100 mM and 1mM (top). Solutions were prepared using mixtures of NaCl:Na_2_SO_4_ (Methods). The V_Cl_ transitions were fitted with single exponentials (red dashed lines), which yielded time constants between 1.5 and 2.6 sec.

The response time of a liquid-junction ISM is usually faster as the width of the organic phase decreases at the tip of the electrode [12,21]. The width for the liquid membrane of our MC3-ISMs was ~300 μm, and we did not explore whether the time constant could be reduced by decreasing the width of the organic phase. In any event, a time response in the order of a few seconds is sufficient to study intracellular [Cl^−^] changes expected to occur in several tens of second to minutes in biological settings.

### The MC3-ISM is insensitive to changes in HCO_3_^−^ or in pH

One of the biggest drawbacks of available Cl^−^ sensing ISMs is their sensitivity to HCO_3_^−^ [7]. The problem posed by HCO_3_^−^ is complex. This ubiquitous intracellular anion is more abundant than Cl^−^ in many cells (e.g. 3-fold higher in mammalian skeletal muscle), and the [HCO_3_^−^] varies greatly depending on pH and pCO_2_ changes. Consequently, for Cl-sensing ISMs that are also sensitive to HCO_3_^−^ there is no simple way to distinguish between real *a_Cl_* changes, and spurious potential shifts caused by changes in [HCO_3_^−^]. Early Cl-sensing ISMs based on anion exchangers had notoriously poor selectivity for Cl^−^ over HCO_3_^−^ [2]. Selectivity for Cl^−^ was greatly improved by using anion-selective carriers, e.g. Mn(III) porphyrins [15], but the physiochemical properties of these first-generation carrier electrodes were unfavorable with instability, drift, and non-Nernstian behavior. The high selectivity for Cl^−^ over HCO_3_^−^ and near-Nernstian behavior of MC3-based solid membrane electrodes [5] showed tremendous promise for developing a Cl-selective liquid membrane ISM suitable for biological applications.

We measured the sensitivity of MC3-ISMs to interference from changes in [HCO_3_^−^] or pH, as a test of suitability for these ISMs to be used for intracellular determination of *a_Cl_*. First, we measured the response of MC3-ISMs in constant 10 mM Cl^−^, a typical value for myoplasmic Cl^−^ in skeletal muscle fibers [1,35], upon which was imposed step changes in HCO_3_^−^ from 1 to 50 mM (Figure 4A), a concentration range far beyond those expected to occur in vivo either in physiological or pathological conditions (i.e. ~13 mM [16]). The potentiometric response of MC3-ISMs was completely insensitive to the presence of HCO_3_^−^ at concentrations as high as 50 mM. This is a remarkable feature of the MC3 Cl-selective carrier, and makes MC3-based ISMs the electrode of choice for biological purposes. Similarly, we tested for pH-dependent alterations of V_Cl_ in response to pH changes from 6.5 to 8.5, which covers the entire physiologic range of expected cytoplasmic values. A three-point Cl^−^ calibration response (1, 10, 100 mM) for an MC3-ISM was not affected by a pH shift from 7.4 to 6.5 or from 7.4 to 8.5 (Figure 4B, *left* and *right*, respectively).

**Fig. 4.**
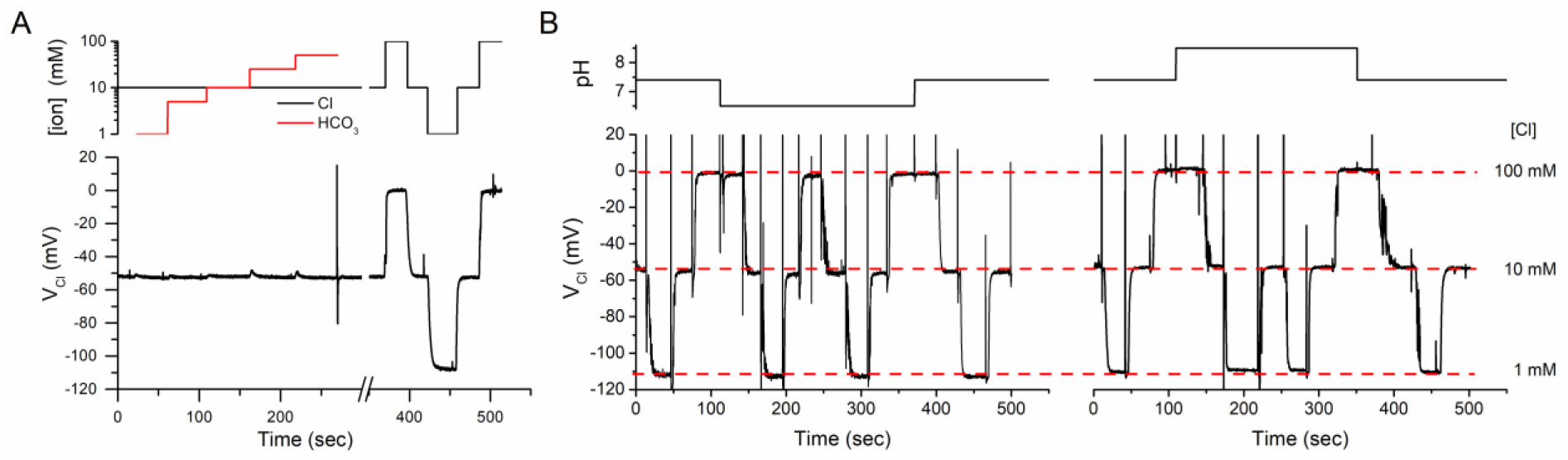
The MC3-ISMs are insensitive to changes in HCO_3_^−^ or pH. (A) The left portion of the V_Cl_ trace (0 to 300 s) shows the MC3-ISM response in 10 mM Cl remained constant, as the [HCO_3_^−^] was varied in step increments from 1 mM to 50 mM. The CO_2_ was nominally 0 in these NaCl:NaHCO_3_ mixtures. The right part of the record (>350 sec) is the response of this same electrode to changes in [Cl^−^] in the absence of HCO_3_^−^. The changes in HCO_3_^−^ and Cl^−^ are depicted in the upper panel by the red and black traces, respectively. (B) The Cl-responsiveness of an MC3-ISM was not affected by pH changes from 6.5 to 8.5 units. The left panel shows the three-point calibration response of V_Cl_ for [Cl^−^] of 1, 10, and 100 mM is identical at pH 7.4 or pH 6.5 (depicted in the upper plot). Similarly, the right panel shows that the Cl^−^ calibration response was not affected by a pH change from 7.4 to 8.5 units. For all panels, the different [Cl^-^ ] were obtained by mixing Cl-based and SO_4_-based solutions (Methods). The “spikes” in the records are due to flux-dependent artifacts during solution switching.

### Chloride selectivity of the MC3-ISM

The selectivity of the MC3-ISM was assessed for Cl^−^ over various physiologically relevant anions including lactate, phosphate, thiocyanate and bicarbonate. The fixed interference method, as described by IUPAC was used [34], wherein the concentration of the test interfering anion was held constant at 10 mM (as a Na^+^ salt), and V_Cl_ was measured in solutions with varying [Cl^−^] between 100 mM and 100 nM. We found that the MC3-ISM response is not significantly affected by the presence of lactate, phosphate, or bicarbonate for [Cl^−^] from 100 mM to values as low as 100 μM (Figure 5). On the other hand, the MC3-ISM was almost insensitive to Cl^−^ in the presence thiocyanate (Figure 5, magenta diamonds). From the data in Figure 5, we calculated the selectivity coefficient, 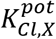, for each test anion (Table 1) which gives a selectivity series: thiocyanate > Cl ≫ bicarbonate > phosphate ≈ lactate. Importantly, the MC3-ISM was 125-fold more sensitive to Cl^−^ than to HCO_3_^−^, which explains the lack of an effect for HCO_3_^−^ in Figure 4 and confirms the MC3-ISM is suitable for measuring intracellular *a_Cl_* without interference from fluctuating levels of HCO_3_^−^.

**Fig. 5.**
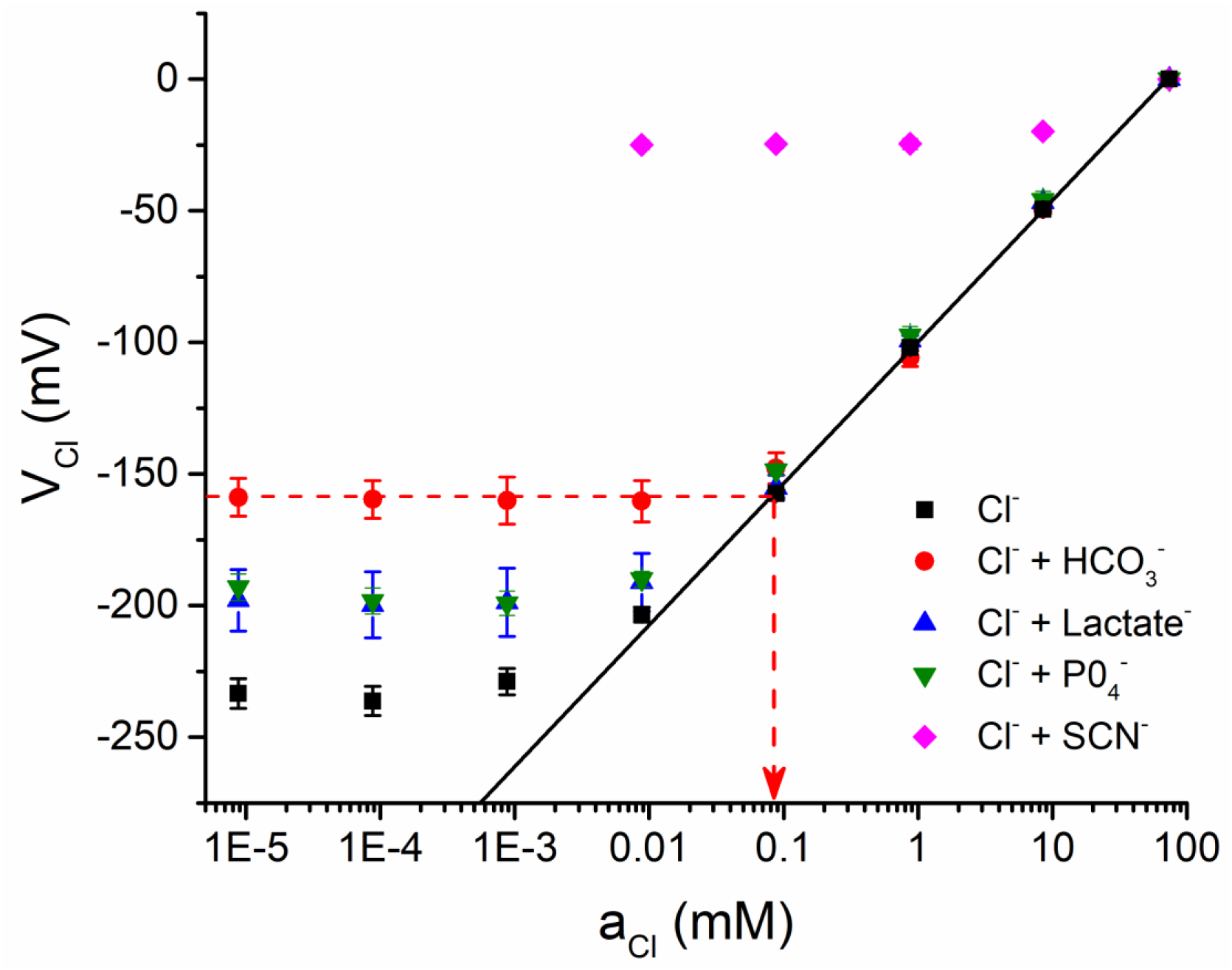
The MC3-ISM is highly selective for Cl^−^. The MC3-ISM response was measured in solutions prepared by serial dilution of pure NaCl with no interfering anions (n = 14, black squares reproduced from Fig. 2) or for the NaCl solutions plus 10 mM of the test anion as Na+ salts of HCO_3_^−^ (n = 3, red circles), lactate (n = 3, blue triangle), PO_4_^−^ (n = 3, green inverted triangle), or SCN-(n = 3, magenta diamond). A linear fit for the pure NaCl data over a range of 0.1 mM to 100 mM solutions shows a Nernstian response (black line). The cut-off *a_Cl_* in the presence of an interfering anion was determined by the intersection of limiting asymptote and the Nernstian response, as shown by the example for HCO_3_^−^ (red dashed line). These values were used to calculate the selectivity coefficient (see Methods).

**Table 1.** Anion selectivity coefficients of the MC3-ISM

| Anion | $a_{Cl}$, (mM) | $K_{Cl,anion}^{pot}$ | $\log_{10}(K_{Cl,anion}^{pot})$ |
| --- | --- | --- | --- |
| SCN<sup>-</sup> | $26 \pm 2.1$ | $2.8 \pm 0.23$ | $0.45 \pm 0.035$ |
| HCO<sub>3</sub><sup>-</sup> | $0.091 \pm 0.010$ | $0.010 \pm 0.0012$ | $-2.0 \pm 0.052$ |
| Lactate | $0.014 \pm 0.0077$ | $0.0015 \pm 0.00085$ | $-3.0 \pm 0.24$ |
| PO<sub>4</sub><sup>-</sup> | $0.010 \pm 0.0028$ | $0.0011 \pm 0.00031$ | $-3.0 \pm 0.15$ |

### Chloride channel blockers and inhibitors of Cl^−^ co-transporters have little effect on the MC3-ISM response

Chloride-selective electrodes are often sensitive to blockers of Cl^−^ channels or to inhibitors of Cl^−^ co-transporters [1,22,8]. We assessed the MC3-ISM sensitivity for two Cl-agents commonly used in muscle research, the ClC-1 blocker 9-ACA and the NKCC1 inhibitor bumetanide (BMT). Our test for interference was based on detecting a distortion of the Nernstian behavior of an MC3-ISM over an operating range of 1 mM to 100 mM [Cl^−^], while the Cl-agents were added at concentrations typically used for studies of muscle. We found that the MC3-ISM is almost insensitive to 9-ACA (Figure 6A and 6C), with a very small effect (reduced V_Cl_ response) in 200 μM 9-ACA when [Cl^−^] was 5 mM or lower. Studies in skeletal muscle that are designed to eliminate the ClC-1 conductance typically use 100 μM 9-ACA [24]. With this concentration of blocker, the V_Cl_ response remained Nernstian down to a [Cl^−^] of 2.5 mM (the lowest expected physiologically), demonstrating that MC3-ISMs can be used to measure intracellular *a_Cl_* in studies with 9-ACA. The MC3-ISM response was insensitive to BMT (Figure 6B and 6D). At 20 μM, a concentration in 10-fold excess of that used for complete inhibition of NKCC1, there was no detectable distortion of the MC3-ISM Nernstian behavior over a [Cl^−^] range from 1 mM to 100 mM.

**Fig. 6.**
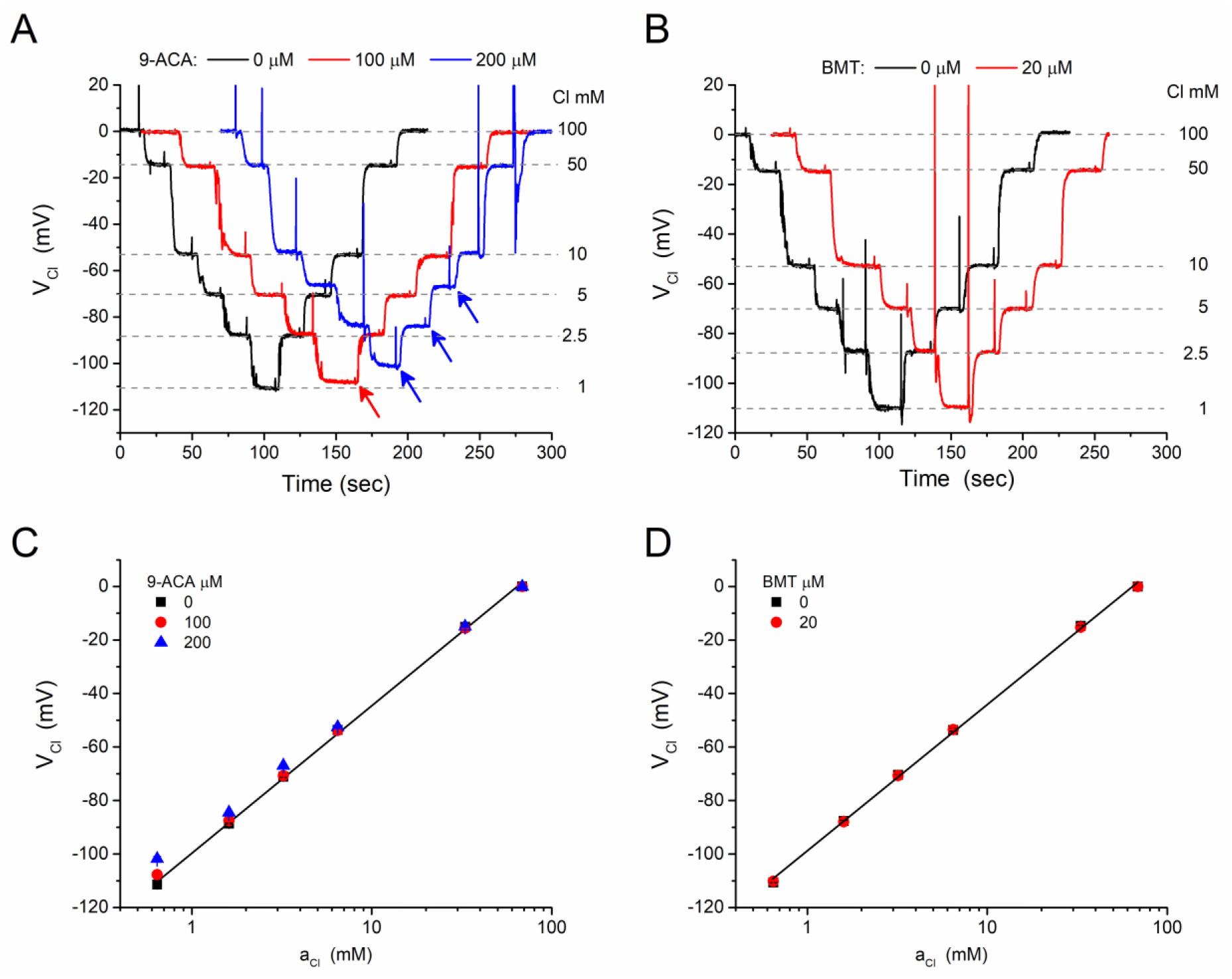
The MC3-ISM response is minimally affected by 9-ACA and is not altered by bumetanide. (A) Continuous records of the MCS-ISM potential (V_Cl_) in response to step changes of [Cl^−^] (1, 2.5, 5, 10, 50, 100 mM) in the absence (black trace) or presence of 100 μM (red trace) and 200 μM (blue trace) 9-ACA. Records for each 9-ACA concentration were obtained consecutively for the same MC3-ISM, but were shifted in time and superimposed to facilitate a visual comparison. Arrows indicate detectable differences from the control response in the absence of 9-ACA. Similar responses were obtained for four different MC3-ISMs. (B) Addition of 20 μM bumetanide (BMT) did not detectably alter the V_Cl_ response of an MC3-ISM. The test protocol and display of the continuous data were the same as in (A). Similar responses were obtained for four different MC3-ISMs. (C, D) Calibration plots of V_Cl_ as a function of *a_Cl_* shows that a modest shift of V_Cl_ was detectable only for 200 μM 9-ACA and at the lowest concentration test solution with 0.1 mM Cl^−^. Symbols represent the mean ± SEM for 4 different MC3-ISMs, with the error bars being obscured by the size of the symbols.

### Intracellular Cl activity, 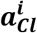, measured with an MC3-ISM

We verified the performance of the MCS-ISM as a sensor of intracellular Cl^−^, by recording 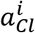 in a skeletal muscle fiber isolated from the flexor digitorum brevis of the mouse. The fiber was impaled with a standard reference electrode to measure the resting potential (V_m_ Fig. 7, upper panel) and also with a Cl-selective MC3-ISM (V_ISM_) which reports the combined effect of V_m_ and V_Cl_. The offset for V_ISM_ = 0 mV was set in a calibration solution with 100 mM [Cl^−^]. The difference between these two signals, V_Cl_ = V_m_ - V_ISM_ (Figure 7, lower panel) is proportional to 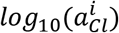.

**Fig. 7.**
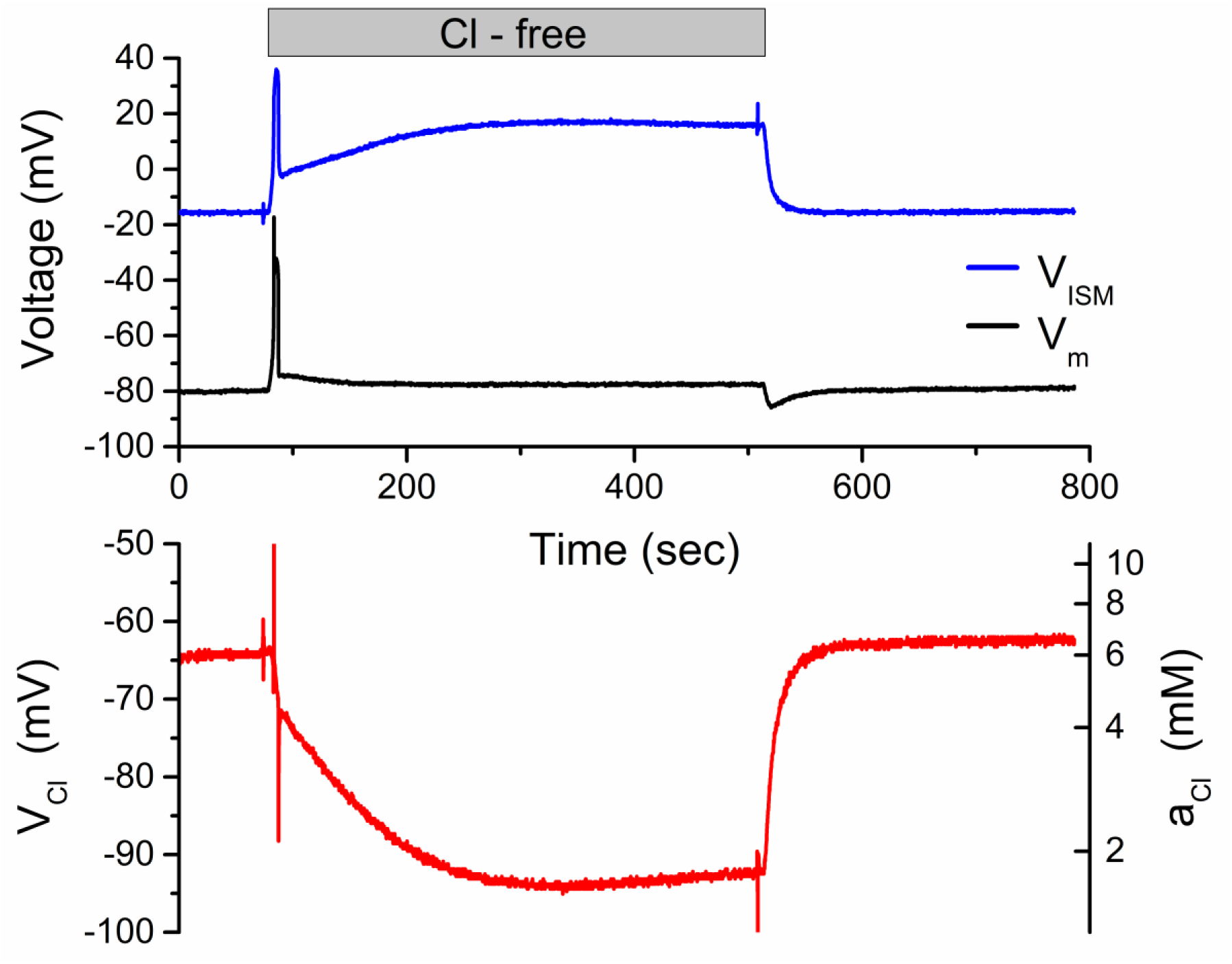
Myoplasmic 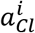 measured with an MC3-ISM. Simultaneous intracellular recordings of membrane potential (V_m_, black trace) and the MC3-ISM response (V_ISM_, blue trace) were obtained by impaling a dissociated fiber from the flexor digitorum brevis muscle (upper panel). Digital subtraction (V_m_ – V_ISM_) was performed to calculate the red trace in the lower panel, representing both linear changes of V_Cl_ (left ordinate) as a function of time and logarithmic changes in 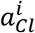 as a function of time (right ordinate). At 80 s, the bath was exchanged to Cl-free Tyrode solution, and then back to normal Tyrode (150 mM Cl) at 500 s.

In control Tyrode solution (140 mM Cl^−^), the resting membrane potential was stable (V_m_ = −80 mV), as was V_Cl_ = −65 mV, corresponding to 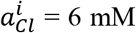. Upon switching the perfusion to SO_4_-Tyrode (nominally zero Cl^−^) the intracellular 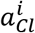 exponentially fell to about 1.5 mM because of Cl^−^ efflux through the high resting conductance of ClC-1 channels. These changes are fully reversible, as shown by the recovery of 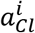 when the perfusion with Cl-Tyrode was resumed at 500 sec in Figure 7.

The transient response of V_m_ is also consistent with decrease of 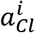, followed by recovery [14]. At the onset of Cl-free Tyrode solution (Figure 7, 80 sec), V_m_ depolarized because the equilibrium potential for Cl^−^ suddenly shifted from about −77 mV to a positive value. Intracellular Cl^−^ content then declined over the next 30 s, which shifted E_Cl_ to more negative potentials and caused V_m_ to relax back toward −80 mV. When extracellular Cl^−^ is suddenly restored (Figure 7, 500 sec), V_m_ had a hyperpolarizing undershoot because intracellular 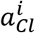 was now lower than baseline. As 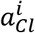 increased back to the normal basal level, V_m_ relaxes to the initial value of −80 mV.

## DISCUSSION

The development of a highly-selective and sensitive chloride sensor in polymeric membrane electrodes, based on the preorganized macrocyclic Lewis acid MC-3 as the anion carrier [5], was a major technological advance. Prior electrode designs using anion exchangers (e.g. TBT or TDMAC) lacked selectivity, which in these electrodes was dominated by anion lipophilicity (Hofmeister series). Improved selectivity was achieved with metalloporphyrin carriers (e.g. Mn(III), chloride ionophore I Sigma-Aldrich) or Hg(II)-organics (chloride ionophore II, Sigma-Aldrich), but these compounds were not suitable for liquid membrane ion-selective microelectrodes because of interference from HCO_3_^−^, electrical instability with drift and non-Nernstian responses, or poor solubility in NPOE. In contrast, MC-3 is chemically stable, and MC-3 based polymeric membrane electrodes have more consistent Nernstian responses, high selectivity for Cl^−^, and faster settling times of a few seconds.

We now show that the advantages of MC-3 based polymeric membrane electrodes can be realized in a liquid membrane microelectrode form factor. Following from the design of the MC-3 electrodes in PVC membranes [5], we used NPOE as the liquid organic phase in our ISMs and no other plasticizers were tested. Just as with the PVC membrane electrodes, the potentiometric response to Cl^−^ with MC-3 alone was sub-Nernstian (Figure 1), and doping the liquid membrane with a lipophilic cationic additive (2.5% TDMAC) markedly increased the ISM sensitivity to near-Nernstian behavior (54.3 mV/decade). Increased amounts of cationic additive in PVC membrane electrodes extended the linear range for high *a_Cl_* to about 100 mM, but with a cost of slightly reduced selectivity for Cl^−^ over other anions [5]. Since our ISMs already had linear responses up to the highest concentration of the calibration solution (nominally 100 mM mixture, calculated *a_Cl_* = 68.5 mM), we did not test higher amounts of additive. Moreover, in a physiological context the expected intracellular *a_Cl_* is in the range of 4 to 40 mM.

The potentiometric responses of MC-3 ISMs were remarkably consistent and stable over long recording periods. Except for the occasional failure in the fabrication of an ISM (e.g. air bubble at the liquid membrane / aqueous electrolyte interface), the variability was exceptionally small in the slope of the linear response range (0.1 mM to 100 mM). A sample of 24 MC-3 ISMs with an average slope of 54.6 mV/decade had a standard deviation of only 0.9 mV/decade and a range of 52.9 to 56.0 mV/decade. By comparison, a sample of 62 ISMs constructed with the “improved” Corning 477913 chloride ionophore was reported to have a mean slope of 52.8 mV/decade with a standard deviation of 5.1 mV/decade and the range of values extended from 35 to 64 mV/decade [6]. The potentiometric response of MC-3 ISMs was stable with no significant drift over ten minutes or more, as shown by examples in several Figures (2–4, 6). Moreover, the intracellular response recorded from a skeletal muscle fiber was also stable, as shown by the return to the baseline value after recovery from a 10 min exposure to Cl-free conditions (Figure 7).

The slow electrical response time of Cl-selective ISMs has been a limiting factor for use of some ionophores. The problem arises from the high resistivity of immiscible liquid membranes with neutral ion carriers in combination with the capacitance of the glass pipette. While reducing the ISM tip resistance with cationic additives, choice of the ionophore, and optimization of the liquid membrane thickness may be used to improve performance, the 10% to 90% response time was reported to be as long as 41 sec for Mn(III) porphyrin Cl-selective ISMs [17]. For macroscopic MC-3 electrodes in PVC membranes, a more favorable rise time of about 10 sec was observed [5]. In our MC-3 ISMs the 10% to 90% response time was about 5 sec (Figure 3), which is sufficient to monitor changes of intracellular *a_Cl_* in muscle fibers that typically occur over tens of seconds to minutes.

The most important advantage of MC-3 ISMs is the high selectivity for Cl over other anions, especially HCO_3_^−^. Historically, the Cl-ISMs used in several landmark papers showing intracellular *a_Cl_* in muscle and heart is higher than expected from passive electrodiffusion were based on proprietary Cl-ionophores from Corning (477315 and later 477913) that had substantial interference from HCO_3_^−^ [1,7,35]. This limitation was compounded by the composition of the intracellular milieu, where *a_Cl_* is typically 4 mM, and HCO_3_^−^ is more abundant at 14 to 18 mM. With a Corning 477913 ISM, the selectivity for Cl over HCO_3_^−^ was about 8:1, corresponding to a selectivity coefficient 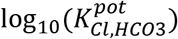 of −0.9 [17]. With this degree of HCO_3_^−^ interference, a true *a_Cl_* of 4 mM in combination with an *a*_*HCO*3_ of 15 mM would cause a 9.3 mV error in the potentiometric value of the Cl-ISM, corresponding to an overestimate of *a_Cl_* as 5.9 mM. In contrast, the high Cl-selectivity we observed for an MC3-ISM, 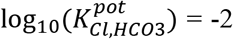, would result in only a 0.9 mV error with an estimated value of *a_Cl_* = 4.14 mM. Insensitivity to HCO_3_^−^ (Figure 4) is also important because intracellular pH often changes or is intentionally manipulated during a study. The accompanying fluctuations of intracellular *a*_*HCO*3_ would render the potentiometric response uninterpretable for a poorly selective Cl-ISM. Sensitivity to pH is also a limitation for genetically encoded Cl sensors based on YFP variants, which has led to the development of ratiometric dual-sensor fluorescent proteins (ClopHensor) to monitor both [Cl] and pH simultaneously [4].

Another selectivity-based limitation of earlier Cl-ISMs was interference from drugs used to block Cl^−^ channels (e.g. 9-ACA) or to inhibit Cl^−^ exchangers and cotransporters (e.g. SITS, bumetanide, furosemide). These compounds are often used in studies of Cl transport, where drug-dependent changes in extracellular and intracellular Cl activity are measured with ISMs. These inhibitors have high lipid solubility and are anions at physiologic pH, and so not surprisingly, substantial anion interference occurs for ISMs using poorly selective ionophores such as Corning 477413. For example, furosemide at 100 μM caused nonlinear potentiometric responses for *a_Cl_* ≤ 30 mM, with a corresponding selectivity coefficient of 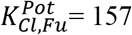 [8]. In contrast, the potentiometric response of the MC3-ISM over the entire range of test solutions ([Cl] from 1 to 100 mM) was not affected by 20 μM bumetanide (Figure 6), which is ten times the IC_50_ to inhibit the Na-K-2Cl 1 cotransporter. For the ClC-1 channel blocker, we detectable interference in 200 μM 9-ACA when [Cl] < 10 mM (Figure 6A). Studies in skeletal muscle typically use 100 μM 9-ACA, and at this lower concentration we detected a change in the MC3-ISM response only for [Cl] < 2.5 mM. Overall, the MC3-ISM is highly selective for Cl with a linear Nernstian response over the biologically meaningful operating range of 1 to 100 mM, with no interference from endogenous anions (10 mM HCO_3_^−^, PO_4_^−^, or lactate) or bumetanide and very modest effects in high-dose 9-ACA well above the IC_50_.

Finally, we demonstrate that the MC3-ISM reliably measures *a_Cl_* in a biological context, the intracellular activity in an isolated skeletal muscle fiber (Figure 7). The response to an external Cl-free challenge is a stringent test of the MC3-ISM because the resulting decrease of intracellular *a_Cl_* would reveal effects from interfering intracellular anions or other sources of distortion as an attenuation of the change in the potentiometric signal. The response in Figure 7 clearly shows a large response for V_Cl_ = −(V_MC3-ISM_ – V_ref_), corresponding to a decrease of intracellular *a_Cl_* to less than 2 mM. The record in Figure 7 also shows the high quality of the signal-to-noise for measuring intracellular *a_Cl_*, as well as the stability of the response with no significant drift and a return to the same baseline when external Cl is restored 10 minutes later.

## ACKNOWLEDGMENTS

This work was supported by grants AR63182, AR42703 (NIAMS/NIH) and RG-381149 (MDA) to SC, and R21-AR067422 (NIH) to MD. AMS thanks the UCLA Department of Chemistry and Biochemistry for start-up funds and 3M for a Non-Tenured Faculty Award, the Alfred P. Sloan Foundation for a Fellowship in Chemistry, Research Corporation for Science Advancement (RCSA) for a Cottrell Scholar Award, and the NIH for a Maximizing Investigators Research Award (MIRA, R35GM124746). RMD was supported by a National Defense Science and Engineering Graduate Fellowship.

We thank Dr. J. Lopez Padrino for kindly sharing his knowhow on the fabrication of ISM’s. Compound-3 was a kind gift from Dr. Dan Yang (The University of Hong Kong, P.R. China); Compound-4H was a kind gift from Dr. Kyu-Sung Jeong (Yonsei University, Korea); and compound-13 and compound-3 were a kind gift from Dr. Anthony Davis (University of Bristol, UK).

## Author Contributions

The experimental design was conceived by MD, SC, and AS. MC3 was synthesized by RD. Measurement of ISM responses were performed by MQ and MD. The paper was written by MD, SC, and AS. All authors reviewed the final manuscript and approved the content.

## Abbreviations

9-ACA: 9-anthracenecarboxylic acid
BMT: bumetanide
ISM: ion selective microelectrode
MC3: [9]mercuracarborand-3
NPOE: 2-nitrophenyl octyl ether
TDMAC: tridodecylmethyl ammonium chloride

